# Independence-based causal discovery analysis reveals statistically non-significant regions to be functionally significant

**DOI:** 10.1101/2025.06.19.660609

**Authors:** Madison Lewis, Shaun Eack, Nicholas Theis, Matcheri S. Keshavan, Konasale M. Prasad

## Abstract

**Background and Hypothesis:** Traditional fMRI analyses often ignore regions that fail to reach statistical significance, assuming they are biologically unimportant. We tested the accuracy of this assumption using causal discovery based-analysis that go beyond associations/correlations to test the causality of one region’s influence over the other. We hypothesized that the network of statistically significant (active network, AN) and non-significant regions (silent network, SN) ***<u>causally</u>*** interact and their features will causally influence psychopathology severity and working memory performance.

**Study Design:** We examined AN and SN during N-BACK task on 25 FHR and 37 controls. Clusters with significantly different activations were juxtaposed to 360 Glasser atlas parcellations. The PC algorithm for causal discovery was implemented. Connectivity of regions with the highest alpha-centrality (HAC) were examined.

**Results:** Seventy-seven Glasser regions were in the AN and the rest were silent nodes. Two regions showed HAC for FHR and HC. Among controls, one HAC region was silent (auditory association cortex) and the other one was active (insula). Among FHR, both were silent nodes (early auditory cortex). These HAC regions in both groups had bidirectional directed edges between each other forming a reciprocal circuit whose edge-weights causally “increased” magical ideation severity.

**Conclusion:** Causal connectivity between SN and AN suggests that the statistically non-significant and significant regions influence each other. Our findings question the merit of ignoring statistically non-significant regions and exclusively including statistically significant regions in the pathophysiological models. Our study suggests that causality analysis should receive greater attention.

## Introduction

Functional brain differences in psychiatric disorders have been examined extensively using functional MRI (fMRI). Traditional approaches compare regional blood oxygenation level dependent (BOLD) signals between groups and report spatially distinct regions that show statistically significant differences^1^. Such regions are considered biologically important for task performance or pathophysiology and are used to build explanatory models. The regions that do not show such differences are not examined further assuming that those regions are biologically not relevant. We hypothesized that many regions that show BOLD responses below a priori statistical threshold may still be biologically significant because the regional BOLD responses are highly correlated with each other. Further, the correlation-based functional networks do not reveal whether activation in one region causally influences other regions with which it is correlated. For these reasons, it is important to examine the significance of functional brain changes using approaches that do not entirely depend on correlations by examining effective connectivity that reveals causal influence of regional BOLD activation on other regions to better elucidate functional networks.

Causal discovery-based network analysis (CDBNA) involves building directed acyclic graphs (DAGs) to examine causal associations between node pairs. Thus, if a directed edge from node *i* to node *j* is present, then, the probability that node *j* showing higher/lower activation may be causally influenced by activation in node *i*. It is important to note that “causality” in this context does not imply a mechanism of causation but rather a direction of interaction. Such an approach can enhance the mechanistic understanding of functional networks that can help characterize targets for novel treatments.

To test this proposition, we examined task fMRI data of a cohort of familial high-risk (FHR) individuals for schizophrenia compared to healthy controls (HC) while performing letter N-back task for working memory. We examined FHR individuals because understanding the neurobiology of familial high-risk for schizophrenia is important for early detection of converters to schizophrenia and other psychopathology. Early detection is important because the long-term social outcomes of schizophrenia have not substantially improved over the last century^2–4^. In this study, we examined FHR individuals who were initially recruited in mid 2000’s and followed up for over 15 years^5–8^. The functional imaging data were collected only at the recent follow up. An advantage in studying this cohort is that the FHR participants were beyond the high-risk period for developing schizophrenia, thus providing a more consummate neurobiology compared to many prior studies that examined younger high-risk individuals when the risk of conversion was continuing to increase. We examined working memory because impairments in working memory have been consistently shown among FHR individuals^9–12^. Prior fMRI studies using working memory task on schizophrenia^9,13^ and FHR^10–12^ reported altered activation in different regions but findings are inconsistent regarding the directionality of change and most regions showing group differences did not significantly correlate with working memory or psychopathology. These data question the usefulness of statistical cutoffs in determining functionally different activation patterns in the brain to improve our understanding of pathophysiology and design new treatments.

We tested three hypotheses: (a) many statistically non-significant (“silent”) regions, considered biologically not relevant, and statistically significant (“active”) regions will causally interact with each other, (b) the combination of undirected and directed networks will provide a more comprehensive picture of causal interaction within and between the network of active (AN) and silent (SN) regions, and (c) not all correlations with network and edge metrics would be “causative” of the changes in cognitive and psychopathological measures. To test our hypotheses, we first analyzed the N-back task fMRI data for regional differences between the groups after applying a statistical threshold, correlated the BOLD response of regions above and below a priori statistical threshold with behavioral measures, examined an undirected functional network, and then applied the CDBNA followed by examining the causal influence of directed network features on N-back performance and severity of psychopathology.

## Methods

### Subject Recruitment

The subject recruitment was described previously^5^. Briefly, FHR were defined as offspring/sibling of a proband with schizophrenia/schizoaffective disorder who had previously participated in our studies^6–8^. Parents/siblings were interviewed on Structured Clinical Interview for DSM-IV (SCID-IV) at initial recruitment. During the follow-up, we interviewed FHR and HC on SCID-IV followed by making “consensus diagnosis” by collating data from SCID-IV, medical records, and collateral information. Subjects with intellectual disability, head injuries with significant loss of consciousness, and neurological diseases were excluded. HC were unrelated to FHR. After fully explaining the experimental procedures, subjects provided informed consents. The University of Pittsburgh IRB approved the study. Psychopathology was assessed using the Chapman scale^14^ and in-scanner working memory accuracy and processing time were obtained from the N-back task.

### Imaging Methods

Imaging methods have been previously described^5^. Briefly, a 7 Tesla Siemens whole-body scanner and custom-built Tic-Tac-Toe radiofrequency head coil which provides improved homogeneity and subject insensitivity compared to standard commercial coils^15–17^ was used to acquire T_1_-weighted MP2RAGE of 348 slices (0.55mm thickness, TR=6.0s). Additionally, N-back task-based fMRI data with 86 slices, 1.5mm thickness, and a TR=2.5s was acquired. N-back task was administered using E-prime with two blocks each of 0-back, 1-back, and 2-back. Subjects responded using a five-finger glove box.

Using FSL 6.0^18^, the fMRI data was motion and bias distortion-corrected, skull-stripped, and visually inspected for optimum segmentation and quality. These images were realigned and unwarped, slice-timing corrected, and coregistered with the T_1_-weighted image using Statistical Parametric Mapping 12 (SPM12) within MATLAB R2019a^19^. Quality of coregistration was manually checked for every subject. T_1_-weighted images were segmented to calculate six tissue priors: gray matter, white matter, cerebrospinal fluid, skull, non-brain tissue, and air. Using the deformation fields created during segmentation, fMRI images were normalized, and registration was checked manually using avg152T1 canonical brain followed by smoothing. Next, the average Montreal Neurological Institute (MNI) brain image was registered to the first volume of the preprocessed N-back data.

### Case-control voxelwise comparisons

We compared voxelwise BOLD signal differences contrasting the 2-back with 0-back timeseries data to examine maximal differences in working memory load. A priori combined intensity and spatial extent threshold was p<0.05 for 8 contiguous voxels. Extent threshold was determined using *3dClustSim*^20^ within Analysis of Functional NeuroImages (AFNI)^21,22^ to ensure the largest noise-only cluster in 15 randomly chosen subjects (∼25% of the sample). The average of largest noise-only cluster size was used as the extent threshold (8 voxels).

Beta weights from clusters containing regions associated with working memory were correlated with accuracy and response time of N-back and prodromal symptom severity using the Chapman scale. Specifically, we included average scores for magical ideation (CHAPAVMI), perceptual aberration (CHAPAPAS), anhedonia (CHAPAVAS), and social anxiety (CHAPAVSAR) along with premorbid adjustment (PMASTOT). Tests were corrected for multiple comparisons using the Bonferroni approach. Voxel clusters associated with N-back performance were identified based on the MNI coordinates of each cluster’s peak intensity voxel location within both the Talairach and Glasser atlases within MNI space. Atlas regions for each cluster were investigated for their association with N-back performance based on published data and information from the Glasser atlas supplemental information^23^.

### Undirected Functional Network

We parcellated the cortex using the Human Connectome Project Multi-Modal (version 1)^23^ (Glasser) atlas into 360 functionally and structurally connected cytoarchitecturally defined regions, allowing for better interpretations of regional functional connectivity. The Glasser atlas in MNI space^23^ was registered to the first volume of the individual N-back data file. Quality of parcellation was checked manually. The down-sampled atlas was then used to extract the entire N-back BOLD signal timeseries from each Glasser region. The regional timeseries was then sectioned into divisions based on N-back load: 0-back and 2-back. The difference between the 2-back and 0-back timeseries was calculated as the 2-back>0-back contrast in group comparisons within SPM12. The 2-back>0-back timeseries was used for each individual in the remainder of this manuscript.

We labeled each Glasser parcel as active, silent (as stated), or noise. The statistical threshold was applied to the comparisons on SPM12. Although an extent threshold of 8 voxels was used previously to filter *clusters* below a priori statistical threshold stated above, a more conservative approach was used as our *parcel threshold* to identify Glasser regions to minimize the noise. The noise-reducing 12 voxel parcel threshold was chosen because the point spread function (PSF) of BOLD signal using 7T is 0.8-1.2mm depending on cortical depth with isotropic voxel size=0.5mm.^24^ At a maximum, the PSF was 2.4x the size of the voxel. To estimate PSF, we multiplied the voxel size (2.5mm isotropic) by 2.4 for a PSF=6mm. Using the published evidence^24^, we estimated this number in all three directions (PSF=216mm^3^). A region with less than 12 voxels in a cluster was considered noise and not included in the analyses **(Figure 1B)**.

**Figure 1:**
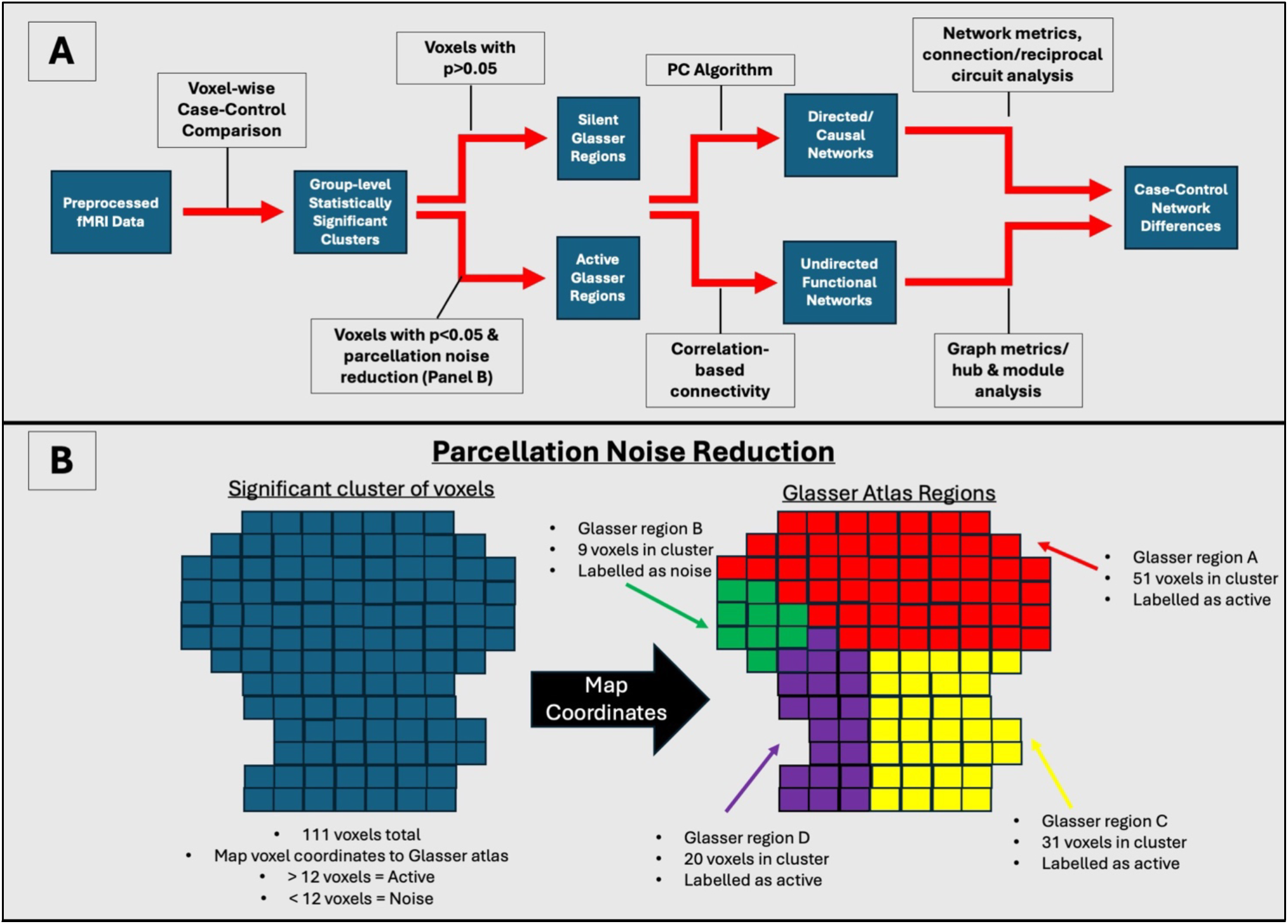
A) Flowchart of analytical approach. The raw images are first preprocessed using FSL and SPM. These images are then parcellated using the Glasser atlas. The outputs of preprocessing are loaded into SPM12 for case-control voxelwise BOLD signal comparisons. B. Parcellation Noise Reduction. All significant clusters from the case-control comparisons were mapped to the Glasser atlas. Each block in the figure is a 2D representation of a voxel. Glasser regions are labelled as active regions (red, yellow, and purple) if 12 or more voxels are within the cluster. If less than 12 voxels are in the cluster (green), that region is labelled as noise.

We tested the relationship of average BOLD signals for each active and silent regions on the Glasser atlas with psychopathology (CHAPAVMI, CHAPAPAS, CHAPAVAS, CHAPAVSAR, and PMASTOT) and working memory scores (N-back accuracy and response time) using partial correlation tests. False discovery rate (FDR)-corrected significances are reported. These correlations are different from the correlation of β weights described above.

For each subject, a full network of 360 Glasser regions was built using the Pearson correlations between regions in the 2-back>0-back contrast. We similarly built one network of only active regions and another of only silent regions for each subject. Each network was absolutized and binarized using the percolation threshold^25^.

Nodal (betweenness centrality, eigenvector centrality, clustering coefficient, and degree) and global (modularity, pathlength, and assortativity) graph metrics were calculated on all networks to understand network integration, segregation, centrality, and community structure^26^. T-tests were used to compare network metrics between groups and ANCOVA to compare these metrics between AN and SN covarying for network sizes. Multiple testing was corrected using the Bonferroni approach. Significant graph measures were correlated with psychopathology and working memory scores using partial correlation followed by correction for multiple testing using FDR approach. Community structure was investigated for averaged group-level networks and hubs were identified when nodes within the modules showed regional degree, betweenness centrality, or eigenvector centrality >2 standard deviations above the mean^26,27^. The top 5% of hubs across the entire group were considered group-level hubs.

### PC Algorithm

We used the Peter Spirtes and Clark Glymour (PC) algorithm to estimate DAG and examine causal connections between and within the regions in AN and SN^28,29^. The PC algorithm uses conditional independence to remove edges in the complete graph and then determine direction of the edge using v-structures and orientation propagation. V-structures use conditional independence to help distinguish different possible causal structures and orientation propogation orients the edges based on the presence/absence of paths in the graph^30^. The advantages with PC algorithm include having no limits on the number of regions-of-interst, overlaps with structural connections better, and measures instant causality without the need for lagged information.

For each subject, the average 2-back>0-back timeseries for every active and silent region was used as an input to the PC algorithm while excluding noise regions. Using the *pcalg* package version 2.7-8 in R version 4.2.2, the PC algorithm was used to make a directed network for FHR and HC separately. Connections were considered significant if α<0.01 for the conditional independence tests. Directed edges between active-to-active, active-to-silent, silent-to-active, or silent-to-silent regions were identified and grouped as commissural (interhemispheric) and associational (within the hemisphere). Network metrics specific to DAGs (α-centrality, assortativity, betweenness, degree, density, and reciprocity) were calculated using the *igraph* network analysis package^31^. The BOLD response from regions with highest α-centrality, which are considered the most important regions for network communication, were correlated with psychopathology severity and task performance using partial correlation tests.

Next we investigated the presence of reciprocal circuits that were defined as a subnetwork with ≥2 nodes where the directed edges were present in both directions, i.e. *i*®*j* and *j*®*i*. We categorized these circuits into active-active, silent-silent, and silent-active based on the types of regions that were bidirectionally connected. Because we were specifically interested in the active-to-slient region interactions, we compared the active-silent circuits between groups and correlated the α-centrality of the regions involved in the circuits as well as the edge weights of the circuits to psychopathology and task-performance measures. Followed by this, we tested whether significant correlations of psychopathology and working memory performance with reciprocal circuit metrics were causative using the PC algorithm. The results are shown as a heat map to highlight the differences between correlations and causal influence.

First, we report the results of BOLD response differences between FHR and HC using an intensity and extent threshold corrected statistical cut-off and correlations with behavioral measures followed by results of undirected graph metrics including differences in AN and SN. Next, the results of CDBNA for the network and for behavioral scores are presented.

## Results

### Demographic characteristics

The mean age and sex did not differ between FHR (n=25; 28.70±3.78 years; 54% female) and HC (n=37; 30.48±3.54 years; 64% female) (t=1.89 p=0.06; χ^2^=0.61, p=0.44). Parental education was significantly different between HC (14.77±2.24 years) and FHR (13.00±1.68) (t=3.55 p=0.0007) that is typical of this type of sample. The reasons for exclusion of consented subjects from this analysis is in supplemental information.

Among the FHR included, 4 subjects did not have any psychiatric diagnosis. Others had a DSM-IV diagnosis: Schizophrenia (N=1), Schizoaffective Disorder (N=3), Major Depressive Disorder (N=3), Panic Disorder (N=1), and Generalized Anxiety Disorder (N=4), Social Phobia (N=1), Obsessive Compulsive Disorder (N=1), Hypochondriasis (N=2), Alcohol Dependance (N=1), Posttraumatic Stress Disorder (N=3), and Attention Deficit Disorder (N=1).

### Voxelwise BOLD response comparisons and correlations with behavioral measures

SPM-12 analysis showed 22 clusters with FHR>HC contrast and 3 with HC>FHR contrast (**Supplemental Table 1**). Three of the 25 clusters contained regions that are associated with working memory according to literature^32,33^, all from FHR>HC contrast. The peak voxel intensity of these clusters were located to the left ventral anterior cingulate (MNI -2,-2,32), right ventral anterior cingulate (MNI 6,-14,38), and right ventral posterior cingulate (MNI-4,-52,14)^34^. The correlations of cluster β weights with psychophatholgy and N-back scores did not survive Bonferroni correction.

Of the 360 Glasser regions, 77 were active from the 25 voxel clusters, 245 were silent, and 38 regions were considered noise (**Supplemental Figure 1**). Among HC, 1 active region (Right Area PGi) positively correlated with 0-back response time and 29 active regions significantly correlated (18 regions positively and 11 regions negatively) with 2-back response time (**Supplemental Table 3**). 0-back and 2-back response time were also significantly correlated with 51 and 130 silent regions, respectively (**Supplemental Table 4**). These correlations survived FDR correction.

However, BOLD responses in the active and silent regions among FHR did not show significant correlation with N-back scores or psychopathology measures.

## Undirected Functional Network

### Graph Measures

Global and nodal graph measures were not significantly different between FHR and HC. However, nodal graph measures of HC, but not of the FHR, showed significantly higher eigenvector centrality and clustering coefficient in AN compared to SN whereas SN showed higher betweenness centrality and degree compared to AN (Supplemental **Table 5**). Within HC and FHR, we noted significantly lower betweenness centrality but higher eigenvector centrality in the AN compared to the SN (See Supplemental table 6 for average global metrics).

The graph measures, global or nodal, from either the AN or SN were not correlated with the N-back or psychopathology measures after FDR correction.

### Hubs and Modules

Network hubs differed between groups in both the AN and SN. Module assignments were similar between groups for the AN with only 3 regions assigned to different modules. However, the SN module assignments were dissimilar; FHR had 6 modules whereas HC had 3 (See **Supplemental tables 7-9** for details).

## Directed Functional Network via PC Algorithm

### Causal connectivity

FHR full network had 251 directed edges whereas the HC was slightly denser with 299 directed edges. The density of directed edges among HC was significantly higher in silent-to-silent, silent-to-active, and active-to-silent but not in active-to-active compared to FHR (**Table 1A;** χ^2^ for proportions: χ^2^=14.55, df=3, p=0.002). Most edge weights were positive except for 4 negative edge weights for HC. FHR had fewer associational edges (within the same hemisphere) (**Table 1B**). For FHR silent-to-active and active-to-silent connections, only 7% and 5%, respectively, were commissural edges compared to 23% and 11% for HC. Overall distribution of associational and commissural connections between FHR and HC was not significantly different and neither was with each of the silent-to-silent, active-to-active, silent-to-active, and active-to-silent connections. Similarly, distribution of reciprocal circuits **(Table 1C)** was also not significantly different (χ^2^=4.75, df=2, p=0.093).

**Table 1:** Types of Causal Connections: A. Number of directed connections for each group and the active and silent connection interaction. B. Percent of directed connections that are associational, or the percent of connections within the same hemisphere, and commissural, interhemispheric connections. C. Reciprocal circuits in FHR vs HC and the characterization of connection.

| A. Number of Directed Edges between Active and Silent Regions in FHR and HC |  |  |  |  |  |
| --- | --- | --- | --- | --- | --- |
| Type of Connection | Silent to Silent | Active to Active | Silent to Active | Active to Silent | Total |
| **Familial High-Risk** | 161 | 55 | 14 | 21 | 251 |
| **Healthy Control** | 193 | 49 | 22 | 35 | 299 |
| B. Percent of Causal Connections that are Associational/Commissural Connections |  |  |  |  |  |
| Type of Connection | Silent to Silent | Active to Active | Silent to Active | Active to Silent | Total |
| **Familial High-Risk** | 88%/12% | 87%/13% | 83%/7% | 95%/5% | 89%/11% |
| **Healthy Control** | 85%/15% | 92%/8% | 77%/23% | 89%/11% | 86%/14% |
| C. Number of Reciprocal Circuits Between Active and Silent Regions in FHR and HC |  |  |  |  |  |
| Type of Connection | Silent to Silent | Active to Active | Active to Silent | Total |  |
| **Familial High-Risk** | 46 | 24 | 10 | 80 |  |
| **Healthy Control** | 44 | 11 | 14 | 69 |  |

### High α-centrality regions and their connections

Since all nodes had α-centrality values, we examined the two nodes in each group with the highest α-centrality. The FHR group’s highest α-centrality regions were both silent and were in the auditory cortex (right Primary Auditory and the right Lateral Belt) whereas one of the HC’s highest α-centrality nodes was active (left Parainsular) and the other was silent (left Area STSv anterior). Both highest α-centrality regions in both groups had bidirectional directed edges between one another with similar edge weights (**Figure 2**). FHR Primary Auditory region had an additional directed edge with the right Medial Belt (silent).

**Figure 2:**
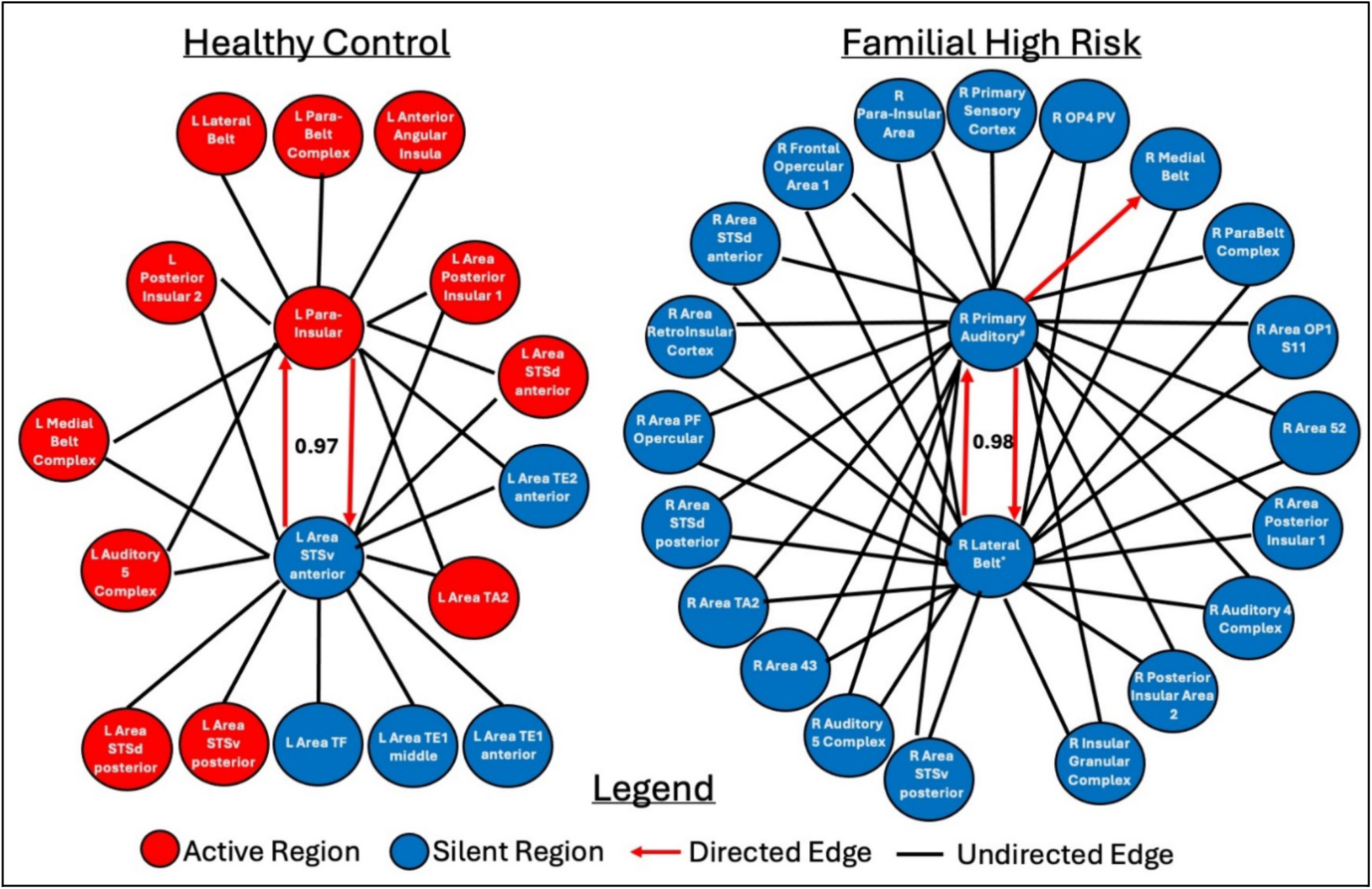
FHR and HC directed and undirected connections with α-centrality regions. Blue indicates a silent region, green indicates an active region, red arrow is a causal connection, black line is an undirected connection

The highest α-centrality nodes of FHR had undirected edges with silent regions only. Both high α-centrality regions had undirected edges with with the same regions except one unique undirected edge (**Figure 2**). Undirected edges of the highest α-centrality regions among HC were to both active and silent regions. Seven of the undirected connections were common between the left Para-Insular and area STSv anterior. All 3 unique connections of the left Para-Insular were to active regions; the left area STSv had 5 unique connections, 2 to active and 3 to silent regions. In both directed and undirected edges of highest α-centrality nodes, connections between active and silent regions were more common among HC whereas FHR showed undirected edges with only the silent regions.

Among HC, BOLD response from the highest α-centrality regions was significantly negatively correlated with 2-back response time (left para-insular, r=–0.52, P_FDR_=0.008; left area STSv anterior, r=–0.52, P_FDR_=0.008) and positively correlated with both 0-back and 2-back response times (right primary auditory, 0-back r=0.62, P_FDR_=0.001; 2-back r=0.67, P_FDR_=0.0003, and right lateral belt, 0-back r=0.64, P_FDR_=0.001; 2-back r=0.71, P_FDR_=0.0001).

The BOLD response of the highest α-centrality nodes of FHR was not significantly correlated with N-back performance after FDR correction.

Correlation of the psychopathology measures did not survive FDR correction for either group.

## Reciprocal Connections

Because the highest α-centrality regions in both groups were connected by reciprocal directed edges, we investigated the reciprocal directed edges in the rest of the network. FHR had 80 reciprocal circuits compared to 69 in HC (**Table 1C**). There were more active-to-active reciprocal connections among FHR and more active-to-silent among HC. Of the active-to-silent reciprocal circuits, most were two-node systems except one three-node system in HC and two in FHR (**Figure 3)**.

**Figure 3:**
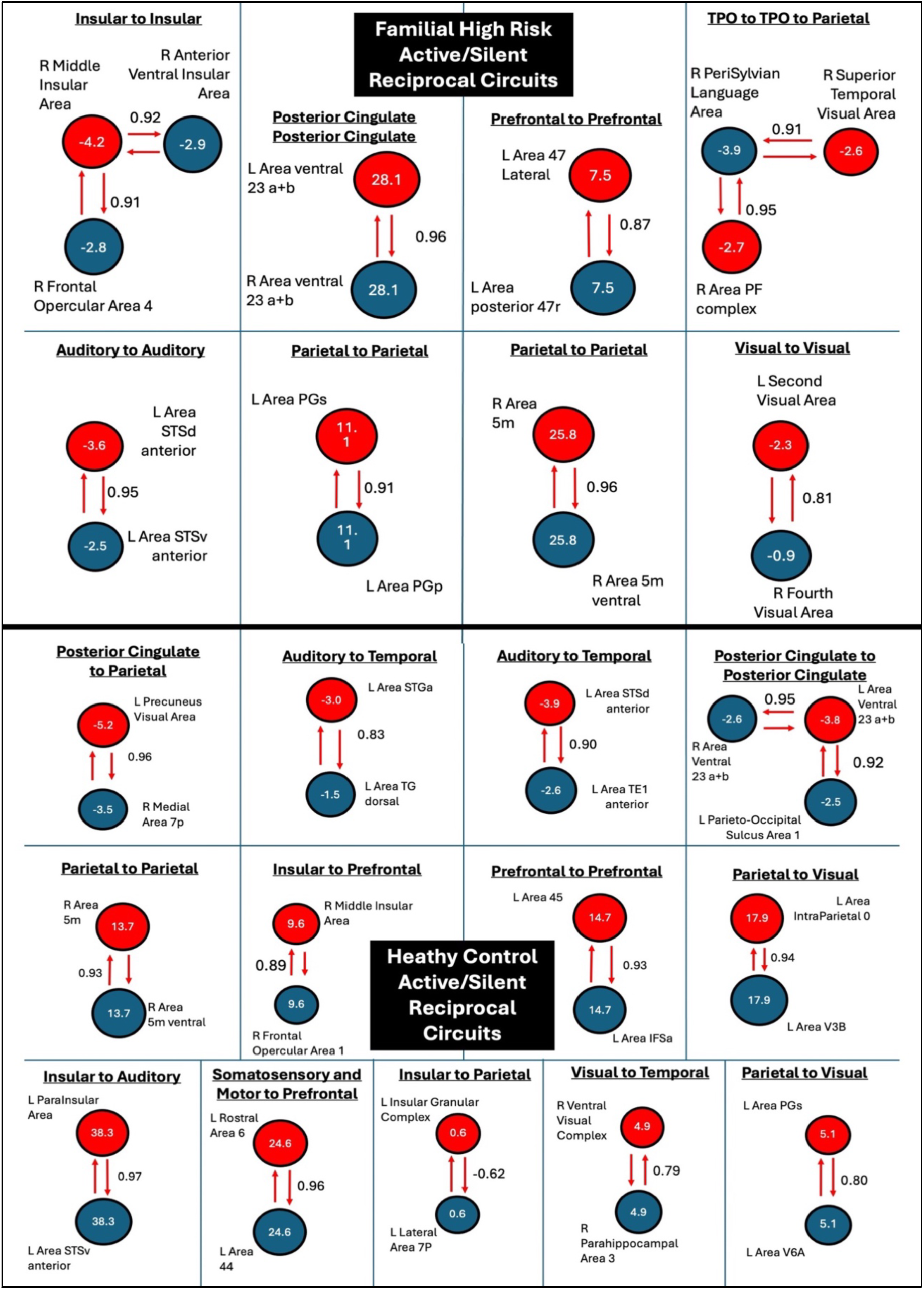
Selected active (red)-to-silent (blue) reciprocal circuits. Alpha-centrality value is shown in the center of each node. Directed connections are indicated using the red arrow and the strength of connection is included. The type of connection according to Glasser sections is also shown above each circuit diagram.

To investigate the association of α-centrality and directed edge weight of reciprocal circuits of active and silent regions with cognition, we first correlated α-centrality and edge weight with the N-back and psychopathology measures. We, then, tested if any variable with significant correlations were also causally influencing one another. We used partial correlation tests controlling for age, sex, and parental education to test correlation of the centrality of each region in the active-to-silent reciprocal circuits and edge weights of each circuit with N-back working memory scores and psychopathology measures using FDR corrections for multiple tests. *Only corrected significances are reported*.

For FHR, the α-centrality of left area ventral 23 a+b (active, Posterior Cingulate Cortex), left Para-Insular area (active, Insular and Frontal Opercular Cortex), and left area STSv anterior (silent, Auditory Association Cortex) were positively correlated with CHAPAVMI (r=0.78, df=15, p=0.008; r=0.73, df=15, p=0.012; r=0.73, df=15, p=0.012) and CHAPAVPAS (r=0.67,df=15, p=0.041; r=0.80, df=15, p=0.004; r=0.78, df=15, p=0.004). The centrality of right area 5m ventral (silent, Paracentral Lobular and Mid Cingulate Cortex) was negatively correlated with 2-back response time (r=–0.74, df=15, p=0.022). For the HC, the centrality of right ventral visual complex (silent, Ventral Stream Visual Cortex) was positively correlated with 2-back response time (r=0.59, df=27, p=0.030).

Only FHR showed significant correlations of edge weights of the reciprocal circuits with working memory scores and psychopathology measures. The reciprocal connections between left area STSd anterior (active, Auditory Association Cortex) and left area TE1 anterior (silent, Lateral Temporal Cortex) as well as the connections of left Para-Insular area (active, Insular and Frontal Opercular Cortex) to left area STSv anterior (silent, Auditory Association Cortex) positively correlated with CHAPAVMI (r=0.89, df=15, p=0.0004; r=0.69 df=15, p=0.028) and CHAPAVPAS (r=0.94 df=15, p=5.1E-7; r=0.75 df=15, p=0.007).

## Behavioral causal analysis by PC algorithm

The PC algorithm consisting of the α-centrality and edge weights of reciprocal circuits showed causal influence on psychopathology severity and working memory performance. Correlations of BOLD response with psychopathology and working memory performance from FHR-HC comparison and graph metrics from undirected network were not tested in the behavioral-causal analysis because the correlations were not significant for FHR.

The directed edge weights of left area STSd anterior-to-left TE1 anterior and of left area STSv anterior-to-left Para-Insular positively causally influenced the CHAPAVMI score (both PC edge weights=0.87) among the FHR. These directed edges also causally influenced the left Para-Insular’s α-centrality (PC edge weight=0.87). The α-centrality of the right ventral visual complex and the centrality of the right area 5m ventral causally connect each other in a reciprocal circuit with edge weight=0.63. Additionally, the directed edge weights in the left area STSd anterior-to-left TE1 anterior and of left area STSV anterior-to-left Para-Insular reciprocally connect to one another with an edge weight=1 (**Figure 4A**). Furthermore, many significant correlations between network metrics and psychopathology scores, some with large effect sizes, for FHR did not show causal influence.

**Figure 4:**
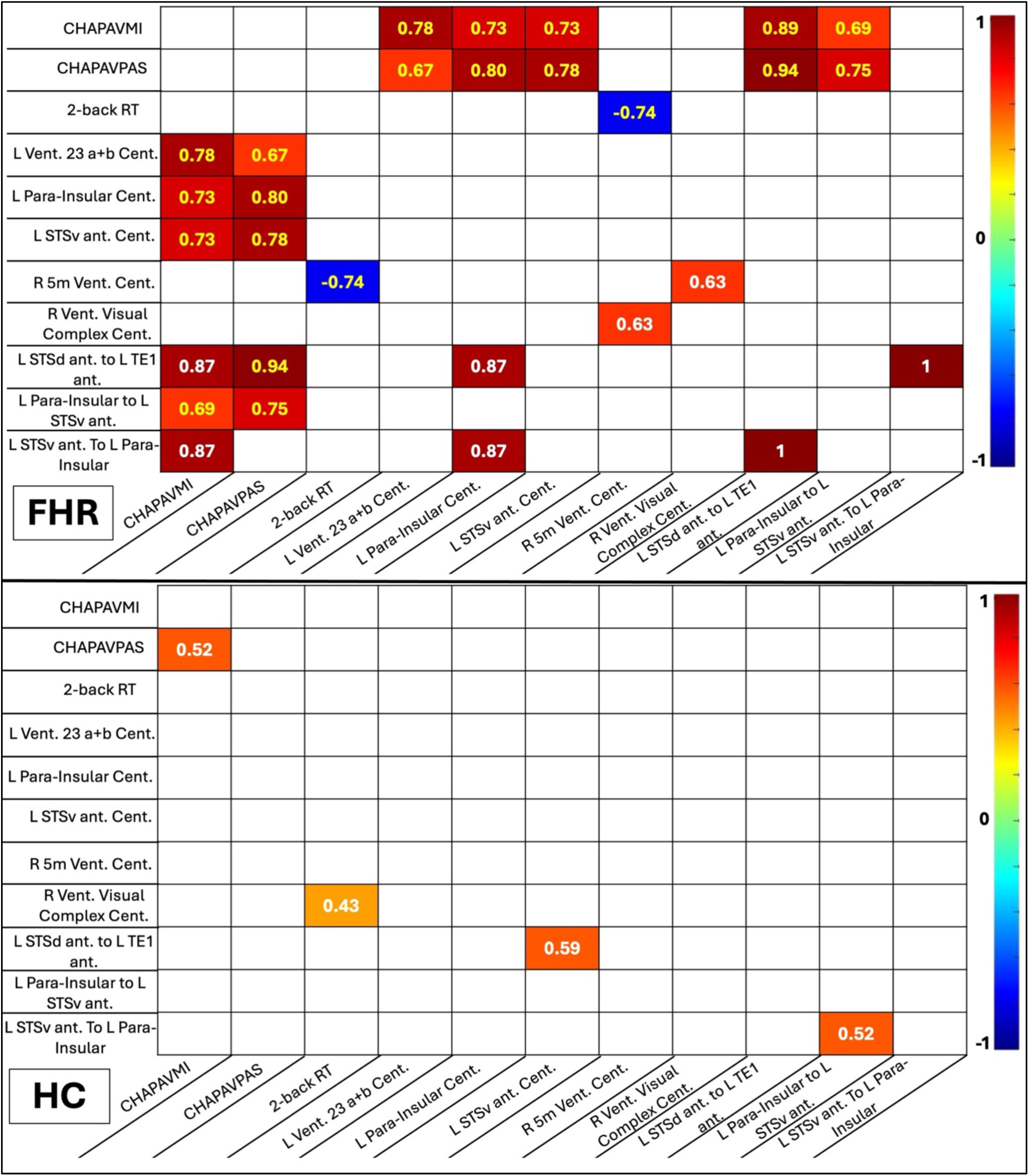
Centrality and edge weights from the reciprocal circuits correlation (yellow font) and causal connectivity (white font) to psychopathology and working memory measures for FHR (top) and HC (bottom). Correlations included in the figure were significant (FDR corrected p<0.05). Color of heatmap indicates the strength of correlation or causal influence (determined using the PC algorithm) using the colorbar on the right. Directed causal influence is read as row influencing the columns. For example in FHR, the connectivity of L STSd ant to L TE1 ant causally influences the CHAPAVMI score with a PC edge weight of 0.87. Abbreviations: RT=response time, Vent=ventral, Cent=centrality, Ant=anterior.

For the HC, four causal influences were identified (**Figure 4B**). They were: the influence the edge weights of left area STSd anterior-to-left area TE1 anterior on the centrality of left area STSv anterior (PC edge weight=0.59), the positive causal influence of CHAPAVPAS on CHAPAVMI, edge weights of left area STSv anterior-to-left parainsular influence on the edge weights of left parainsular-to-left area STSv anterior both with a PC edge weight=0.52, and the centrality of right ventral visual complex causal connection on 2-back response time (PC edge weight=0.43).

## Discussion

The primary motivation for our investigation was to understand whether the statistically non-significant regions (“silent”) that are not considered in pathophysiological models or correlations with cognitive and psychopathological measures are nonetheless causally important. To accomplish that, we examined functional networks of regions that show BOLD signal differences among FHR and HC subjects both above and below a reasonably stringent *a priori* statistical threshold for causal relations using an innovative causal discovery-based analysis. Notably, the silent regions causally influenced active (statistically significant) regions in both groups. Such causal interactions between nodes in AN and SN were more enriched among HC compared to FHR that showed more interactions among nodes within the SN. Such causal relations influenced psychopathology severity and cognitive performance in both groups. These findings suggest that activation of brain regions are more heterogenous and traditional statistical threshold may exclude causally important regions in both groups. The conventional analysis of correlating staticially significant voxel clusters with working memory or severity of psychopathology did not show significant correlations in both groups. But, the Glasser regions underlying the clusters showed such correlations in HC but not in FHR after correcting for multiple testing and excluding noise regions. Thus, the pathophysiological models built using traditional approaches may not capture the neurobiological underpinnings fully as demonstrated by causal influence of directed network features on working memory and psychopathology severity. Further, many correlations with psychopathology and cognitive performance, even those with large effect sizes, were not causally influencing the behavioral measures. Thus, our study suggests that analyses relying entirely on statistical cutoffs provide limited information about functional significance of regions and their influence on behavioral measures whereas causality-based analyses may reveal clinically more meaningful results for building better pathophysiological models and develop novel treatments.

Prior studies reveal that regional BOLD responses are domain-specific^35^, therapies that improve long-term functional outcome enhance regional activations and alter functional connectivity^36^, has revealed important differences between patients and HC, and showed a high degree of accuracy in predicting case-control status^37–40^. However, correlation-based functional connectivity consists of undirected edges that do not provide directionality of interaction, and hence causation. To investigate causal relations, we examined whole-brain directed connectivity, using an innovative and comprehensive CDBNA. Several CDBNA methods exist for fMRI timeseries data to calculate the direction of correlations such as Greedy Equivalence Search (GES), Group iterative multiple model estimation (GIMME), and PC algorithm^41^. GES is a forward selection and backward elimination technique based on the Bayes Information Criterion that requires a large number of timepoints. The length of time-series in a typical fMRI study may be unreliable for GES. A modified version of GES that utilizes information across individuals (Independent Multiple-Sample GES, IMaGES) may make this approach more reliable. Dynamic Causal Modeling considers nonlinear relationships that happen during a task or scan but cannot handle whole-brain connectivity. The GIMME builds a model using a structural equation modeling framework and is more suitable for individual level networks but is limited to about 25 ROIs. Additionally, this choice would not be suitable for our data since GIMME includes lagged effects and would need a consistent task pattern that our N-back data did not have. Similarly, Granger Causality uses lagged timepoints to determine the causality and therefore would not be a viable choice^41^. To calculate causal networks for our data, we used the PC algorithm of conditional independence which leads to network structure that is improved when compared to the traditional connectivity analysis that relies on correlations to define connections^28^.

We found that the architecture of undirected AN and SN were fundamentally different in HC but not in FHR. HC, but not FHR, showed significantly different nodal metrics between AN and SN. Further, community structure of networks was different between FHR and HC. The hubs of the modules were unique to FHR (the anterior cingulate and medial prefrontal cortex regions) and HC (medial temporal, orbital and polar frontal, and dorsolateral prefrontal cortex regions). Despite such differences, their correlation with behavioral measures were not strong. However, when compared between the groups, the AN and SN architecture of HC were not different from that of the FHR suggesting that both networks were similar in terms of features such as network integration, segregation, centrality, and community structure. Thus, the traditional analytical approaches provide limited information on the network architecture and their relationship to behavioral measures.

Compared to undirected networks, causal networks were different between the two groups while also showing correlations and causal influence on psychopathology severity and working memory scores. FHR group had fewer directed connections compared to HC suggesting that the nodal influence on each other was limited. Additionally, for the causal edges between active and silent regions, the HC showed more connections between different Glasser-defined sections. This suggests that FHR had lesser interaction of active and silent regions which are in different areas of the brain and also have different functions. The groups also had different regions with high α-centrality, an indicator of nodal importance. FHR had two silent regions as their highest α-centrality regions whereas the HC had one silent and one active. This further supports the importance and influence of non-significant regions in both groups but especially FHR.

Our directed network analysis also revealed reciprocal circuits in both groups. These reciprocal circuits may be similar to reverberatory circuits that exhibit prolonged sustained activity, creating a loop where signals continuously re-circulate and are maintained for longer periods. Although reciprocal circuits were once thought to be abnormal, reverberation is now viewed to be a likely mechanism for active maintenance of working memory and motor planning which allow neuronal circuits to sustain persistent activity^42,43^. Abnormal reciprocal circuits are thought to arise from imbalances in excitatory and inhibitory neurotransmission between the connected regions, particularly in the prefrontal cortex, which is crucial for cognitive functions^44^. In schizophrenia, reciprocal circuits potentially contribute to hallucinations and disorganized thinking^45^. In this study, FHR had more reciprocal circuits than HC and differed in the connected regions and in connection type. The FHR network had 10 reciprocal connections between active and silent regions whereas the HC had 12. Although similar in number, the FHR reciprocal circuits were mostly between the same Glasser section, such as auditory to auditory or Insular to Insular connection, but the HC reciprocal circuit involved different sections, such as posterior cingulate to parietal. Possible implications are discussed below. Since this study used fMRI data from a working memory task, the reciprocal circuits in HC networks are likely related to the cognitive demands of the N-back task and the motor planning required to answer using the 5-finger glove. Interestingly, the direction of causality for working memory between FHR and HC were opposite of each other. Furthermore, our examination of active and silent connections to understand how different regions influence the rest of the network suggests that the HC more often used their reciprocal circuits to connect spatially and functionally different regions whereas the FHR group tended to reciprocally connect within the same Glasser section. These variations could be due to the differences in modularity in the undirected networks and the FHR network inefficiency reported previously^27,46,47^. Such insight is possible through causal network analyses but not with traditional correlation-based networks with undirected edges.

Our study also revealed causal influence of networks on psychopathology and working memory. The network metrics in the active and silent reciprocal circuit correlated with FHR N-back performance and psychopathology severity after FDR correction. Other analyses, namely regional BOLD signal correlations from active, silent, or high α-centrality regions showed significant correlations only for HC, therefore we did not include these in the causal-behavioral analysis. In the directed network analysis, both the centrality of regions and edge weights of reciprocal circuits significantly correlated with magical ideation, perceptual aberration, and 2-back response time in FHR. Causal-behavioral analysis showed causal influence on the magical ideation from two separate edges (auditory association to lateral temporal connection and auditory association to insular connection). These causal edges had showed high correlations as well. The highest correlations with psychopathology found between the perceptual aberration and two of the directed edges shown in Figure 4 (r=0.94, 0.80, 0.79) were not causal. These findings show that correlations may or may not suggest causation supporting our findings that causal analyses are required for more informed conclusions that may be clinically useful. Additionally, both edges which were causally linked to psychopathology measures were active-to-silent and silent-to-active connections further emphasizing revisiting a priori statistical cutoffs often used in research since most likely the statistically non-significant regions also play a role in the disorder.

The term causality or cause raises several questions such as exclusivity, preceding events, and consequences of an event caused by another. For example, if a network feature causally influences psychopathology severity, the question is whether the observed network feature is the only variable that causally influenced or in conjunction with other variables, whether another event or variable primed the network feature or psychopathology to be causally linked, and whether greater severity of psychopathology affected the network feature, especially when untreated. This can be further illustrated by another example; a force acting on an object causes it to move but the force caused the object to overcome the inertia that then resulted in the motion and also created momentum for the moving object, consequently requiring less force to move the same object. Thus, intermediary factors and consequent events should be considered to better understand causation. For example, centrality of parainsular region shows causal influence on the edge weight of left STSd-to-left TE1 ant edge weight but parainsular region does not have structural or functional connectivity with these two regions^48^ suggesting that the causation is through some other region or edge that is not examined in this study. In summary, causal analysis in the data collected by investigators may reveal causal influence of one variable on another variable but the entire causal pathway requires further work. We emphasize that the observed causality reported here is mathematically derived that requires biological evidence and that the mathematical causality does not refer to the mechanisms underlying causation that requires separate set of experimets specifically designed to test the observed causation. In any case, causal analyses clearly distinguishes itself from correlational analyses and an important step forward in delineating underlying mechanisms and the design of preventative strategies or treatments.

Our sample is unique and was a follow-up of an FHR cohort who were characterized as adolescents between 2003 and 2010 to investigate diagnostic outcomes and the impact of familial risk on neurobiology regardless of conversion to psychoses during young adulthood. This cohort who are toward the end of the highest risk period for schizophrenia can potentially reveal more established impact of familial high-risk on neurobiology than in younger cohort. In previous analyses on the hippocampal network on the same cohort, we had reported volume loss in the hippocampus, its subregions and other medial temporal lobe structures at baseline using 1.5 Tesla MRI data^49–51^. In a subcohort of the baseline sample, we reported reductions in connectivity in frontal-parietal and cingulo-opercular network during N-back task among FHR^5^ along with hub differences between the FHR and HC in these areas and global network disorganization^5,52^. This study shows that there are both undirected and directed network differences in FHR and HC. Although both undirected and directed network analyses showed differences, the directed network was able to provide insight into which networks causally influenced other regions, psychopathology severity, and working memory performance. Additionally, the directed networks identified reciprocal circuit differences between FHR and HC suggesting that directed networks be used more often in examining the functional connectivity.

A strength of our manuscript is comprehensive investigation of the FHR-HC differences in BOLD response during N-back task, undirected functional connectivity, and causal network analysis followed by causal-behavioral analysis. Our findings are novel and bring to light the limitations of statistical cut-off in identifying the regions-of-interest for disease modeling and understanding the biology of psychopathology and cognitive impairments. Limitations of this study include the relatively small sample size, but due to the nature of this unique follow-up and ultra-high field sample the results are still valuable and acceptable. A larger sample could still have conveyed that same message but could have identified regions/networks with smaller effect sizes. Additionally, we only investigated the top two regions with the highest α-centrality for each group, but expanding the analysis to include more regions could have revealed more information. Instead, our goal was to provide a comprehensive approach to investigate both undirected and directed networks and investigate each element’s contribution to psychopathology. The PC algorithm’s limitation includes the requirement of threshold for conditional independence tests and the potential for low individual level reliability of edge directionality.

Our findings should be replicated using independent samples and other disorders. The findings from our study indicate that statistical significance of BOLD signal differences between case-control groups may eliminate some regions that affect the dysconnectivity seen in FHR. Instead of relying on statistical cutoffs, we suggest investigating both significant and nonsignificant regional impact on both undirected and directed brain connectivity analyses. This may provide a more thorough investigation that could increase the understanding of pathophysiology and therefore more effective treatments.

## Acknowledgements

We want to thank the Pittsburgh Supercomputing Center for providing the computational resources through ACCESS Discover grant BIO200047. Additionally, we would like to thank Dr. Kelvin Lim for his assistance with the causal influence of network measures on psychopathology and working memory scores portion of this manuscript.

