## Supplemental Information for "Independence-based causal discovery analysis reveals statistically non-significant regions to be functionally significant"

**Subject Exclusion Details:**

Of the 58 FHR consented, 29 were not scanned due to inability to fit in the bore of the magnet (N=4), subject living out of area (N=6), metal in the body (N=2), piercings/tattoos (N=5), unable or unwilling (N=6), and refused MRI (N=6). Four subjects were excluded during quality control due to scan quality or insufficient Glasser atlas parcellation. Of the 91 HC consented, 28 subjects had their data acquired after the start of this analysis and therefore were not included. Additionally, 17 were never scanned due to inability to fit in bore (N=3), screen fail (N=5), unable or unwilling (N=3), tattoo (N=1), and refused MRI (N=5). Nine were excluded due to scan quality or insufficient Glasser atlas parcellation.

**Nominal Cluster Correlations with Cognition:**

The correlations of cluster β weights with psychopathology and working-memory scores did not survive Bonferroni correction. Nominally, left ventral anterior cingulate cluster was correlated to 2-back accuracy for both the FHR (r=-0.41; uncorrected p=0.04) and HC (r=0.41; uncorrected p=0.01) groups. HC left ventral anterior cingulate cluster was correlated with both magical ideation (r=-0.350; uncorrected p=0.03) and premorbid adjustment total score (r=-0.35; uncorrected p=0.04). Additionally, HC right ventral anterior cingulate cluster was correlated with CHAPAVMI (r=-0.34; uncorrected p=0.04).

Supplemental Table 1: Significant clusters found in case-control voxel-wise comparisons from SPM12.

| Cluster # | Voxels | Uncorrected p | MNI coordinates | Contrast |
|---|---|---|---|---|
| 1 | 4945 | <0.001 | -54, -58, 28 | FHR>HC |
| 2 | 661 | 0.001 | 2, -40, -14 | FHR>HC |
| 3 | 193 | 0.001 | 6, -20, -6 | FHR>HC |
| 4 | 239 | 0.001 | 10, -82, 28 | FHR>HC |
| 5 | 96 | 0.003 | 52, -46, 26 | FHR>HC |
| 6 | 289 | 0.004 | -2, -40, 52 | FHR>HC |
| 7 | 116 | 0.005 | 48, -28, 30 | FHR>HC |
| 8 | 8 | 0.005 | 10, -38, 6 | FHR>HC |
| 9 | 72 | 0.008 | 2, -74, -4 | FHR>HC |
| 10 | 46 | 0.011 | 46, 20, -10 | FHR>HC |
| 11 | 23 | 0.011 | 14, -42, 6 | FHR>HC |
| 12 | 26 | 0.012 | 4, 28, 44 | FHR>HC |
| 13 | 22 | 0.012 | -38, 10, 44 | FHR>HC |
| 14 | 18 | 0.014 | -2, -22, 30 | FHR>HC |
| 15 | 61 | 0.014 | -2, -2, 32 | FHR>HC |
| 16 | 42 | 0.015 | -6, -22, 8 | FHR>HC |
| 17 | 111 | 0.017 | 12, -60, 28 | FHR>HC |
| 18 | 8 | 0.021 | 28, -20, -22 | FHR>HC |
| 19 | 29 | 0.027 | 4, -88, -2 | FHR>HC |
| 20 | 8 | 0.035 | 6, -14, 38 | FHR>HC |
| 21 | 13 | 0.037 | 2, -56, -12 | FHR>HC |
| 22 | 25 | 0.038 | -4, -52, 14 | FHR>HC |
| 23 | 31 | 0.006 | 26,-56,16 | HC>FHR |
| 24 | 25 | 0.026 | 32,-58,44 | HC>FHR |
| 25 | 8 | 0.040 | 8,-66,-6 | HC>FHR |

Supplemental Table 2: Glasser regions within each SPM cluster. Clusters 1-22 are from FHR>HC contrast and clusters 23-25 are from HC>FHR contrast. Clusters highlighted in green were used to correlate with cognition.

| **Cluster 1 (MNI -54,-58,28)** | **Cluster 2 (MNI 2,-40,-14)** | **Cluster 14 (-2, -22, 30)** |
|---|---|---|
| 'L_Area_PGi' | 'R_PreSubiculum' | 'L_RetroSplenial_Complex' |
| 'L_Area_PF_Complex' | 'R_Ventral_Visual_Complex' | 'R_RetroSplenial_Complex' |
| 'L_Area_OP4-PV' | 'R_ParaHippocampal_Area_1' | **Cluster 15 (-2, -2, 32)** |
| 'L_Area_PFm_Complex' | 'R_Fusiform_Face_Complex' | 'L_Area_33_prime' |
| 'L_Area_TA2' | 'R_Area_TF' | 'R_Area_Posterior_24_prime' |
| 'L_Area_7PC' | **Cluster 3 (MNI 6,-20,-6)** | 'R_Area_33_prime' |
| 'L_Anterior_Agranular_Insula_Complex' | NA | 'L_RetroSplenial_Complex' |
| 'L_Area_PFt' | **Cluster 4 (MNI 10,-82,28)** | 'L_Area_Posterior_24_prime' |
| 'L_Auditory_4_Complex' | 'R_Second_Visual_Area' | 'L_Area_23d' |
| 'L_Area_STSv_posterior' | 'L_Second_Visual_Area' | **Cluster 16 (-6, -22, 8)** |
| 'L_Area_2' | 'R_Third_Visual_Area' | NA |
| 'L_Area_PFcm' | 'R_Sixth_Visual_Area' | **Cluster 17 (12, -60, 28)** |
| 'L_Area_PF_opercular' | 'L_Third_Visual_Area' | 'R_Area_7m' |
| 'L_Area_47l_(47_lateral)' | 'R_Primary_Visual_Cortex' | 'R_Parieto-Occipital_Sulcus_Area_2' |
| 'L_Area_PHT' | 'L_Primary_Visual_Cortex' | 'L_Area_7m' |
| 'L_Primary_Auditory_Cortex' | **Cluster 5 (MNI 52,-46,26)** | 'R_Parieto-Occipital_Sulcus_Area_1' |
| 'L_Area_TemporoParietoOccipital_Junction_2' | 'R_Superior_Temporal_Visual_Area' | **Cluster 18 (28, -20, -22)** |
| 'L_ParaBelt_Complex' | 'R_Area_PGi' | 'R_Hippocampus' |
| 'L_Posterior_Insular_Area_2' | 'R_Area_PFm_Complex' | 'R_ParaHippocampal_Area_1' |
| 'L_Area_Posterior_Insular_1' | 'R_Area_TemporoParietoOccipital_Junction_1' | **Cluster 19 (4, -88, -2)** |
| 'L_Medial_Belt_Complex' | **Cluster 6 (MNI -2,-40,52)** | 'R_Primary_Visual_Cortex' |
| 'L_PeriSylvian_Language_Area' | 'L_Area_31a' | 'L_Primary_Visual_Cortex' |
| 'L_Area_TemporoParietoOccipital_Junction_1' | 'L_PreCuneus_Visual_Area' | **Cluster 20 (6, -14, 38)** |
| 'L_Superior_Temporal_Visual_Area' | 'L_Area_5m_ventral' | 'R_Area_23d' |

| | | |
|---|---|---|
| 'L_Area_IntraParietal_2' | 'R_Area_31a' | 'R_Area_Posterior_24_prime' |
| 'L_Area_STGa' | 'R_Area_5m' | **Cluster 21 (2, -56, -12)** |
| 'L_Auditory_5_Complex' | 'L_Area_5m' | NA |
| 'L_Area_TE1_posterior' | 'R_Area_23c' | **Cluster 22 (-4, -52, 14)** |
| 'L_Primary_Sensory_Cortex' | 'L_Area_23c' | 'L_RetroSplenial_Complex' |
| 'L_Area_STSd_anterior' | 'R_PreCuneus_Visual_Area' | 'L_Area_ventral_23_a+b' |
| 'L_Primary_Motor_Cortex' | 'L_Primary_Motor_Cortex' | **Cluster 23 (26,-56,16)** |
| 'L_Area_3a' | 'L_Dorsal_Area_24d' | 'R_Dorsal_Transitional_Visual_Area' |
| 'L_Area_STSd_posterior' | **Cluster 7 (MNI48,-28,30)** | **Cluster 24 (32,-58,44)** |
| 'L_Area_43' | 'R_Area_PF_Complex' | 'R_Area_Lateral_IntraParietal_dorsal' |
| 'L_Area_IntraParietal_0' | 'R_Area_PFcm' | 'R_Area_IntraParietal_1' |
| 'L_Insular_Granular_Complex' | 'R_Anterior_IntraParietal_Area' | **Cluster 25 (8,-66,-6)** |
| 'L_Lateral_Belt_Complex' | 'R_Area_PFt' | 'R_Second_Visual_Area' |
| 'L_Anterior_IntraParietal_Area' | **Cluster 8 (MNI 10,-38,6)** | 'R_Third_Visual_Area' |
| 'L_Area_FST' | 'R_RetroSplenial_Complex' | |
| 'L_Para-Insular_Area' | **Cluster 9 (MNI 2,-74,-4)** | |
| 'L_Area_OP1-SII' | 'R_Second_Visual_Area' | |
| 'L_Anterior_Ventral_Insular_Area' | 'R_Primary_Visual_Cortex' | |
| 'L_Rostral_Area_6' | 'L_Primary_Visual_Cortex' | |
| 'L_Frontal_Opercular_Area_4' | 'L_Second_Visual_Area' | |
| 'L_Area_PGs' | 'R_Third_Visual_Area' | |
| 'L_Area_45' | **Cluster 10 (MNI 46,20,-10)** | |
| 'L_Medial_Superior_Temporal_Area' | 'R_Middle_Insular_Area' | |
| 'L_Area_1' | 'R_Anterior_Agranular_Insula_Complex' | |
| 'L_RetroInsular_Cortex' | 'R_Area_STGa' | |
| 'L_Middle_Insular_Area' | 'R_Area_47s' | |
| 'L_Piriform_Cortex' | **Cluster 11 (MNI 14,-42,6)** | |
| 'L_Frontal_Opercular_Area_1' | 'R_RetroSplenial_Complex' | |

| | |
|---|---|
| 'L_Area_PH' | 'R_Parieto-Occipital_Sulcus_Area_1' |
| 'L_Area_52' | 'R_PreSubiculum' |
| 'L_Area_47s' | **Cluster 12 (MNI 4,28,44)** |
| 'L_Area_OP2-3-VS' | 'L_Area_8BM' |
| 'L_Area_IntraParietal_1' | 'R_Area_8BM' |
| 'L_Area_47m' | **Cluster 13 (MNI -38,10,44)** |
| 'L_Frontal_Opercular_Area_2' | 'L_Area_8C' |
| 'L_Area_Frontal_Opercular' | 'L_Area_IFJp' |

Supplemental Figure 1: 25 voxel clusters showing significant differences (left) which translates to 77 Glasser atlas regions (right) labelled active (shown in green), 245 silent regions (blue), and 38 noise regions (gray).

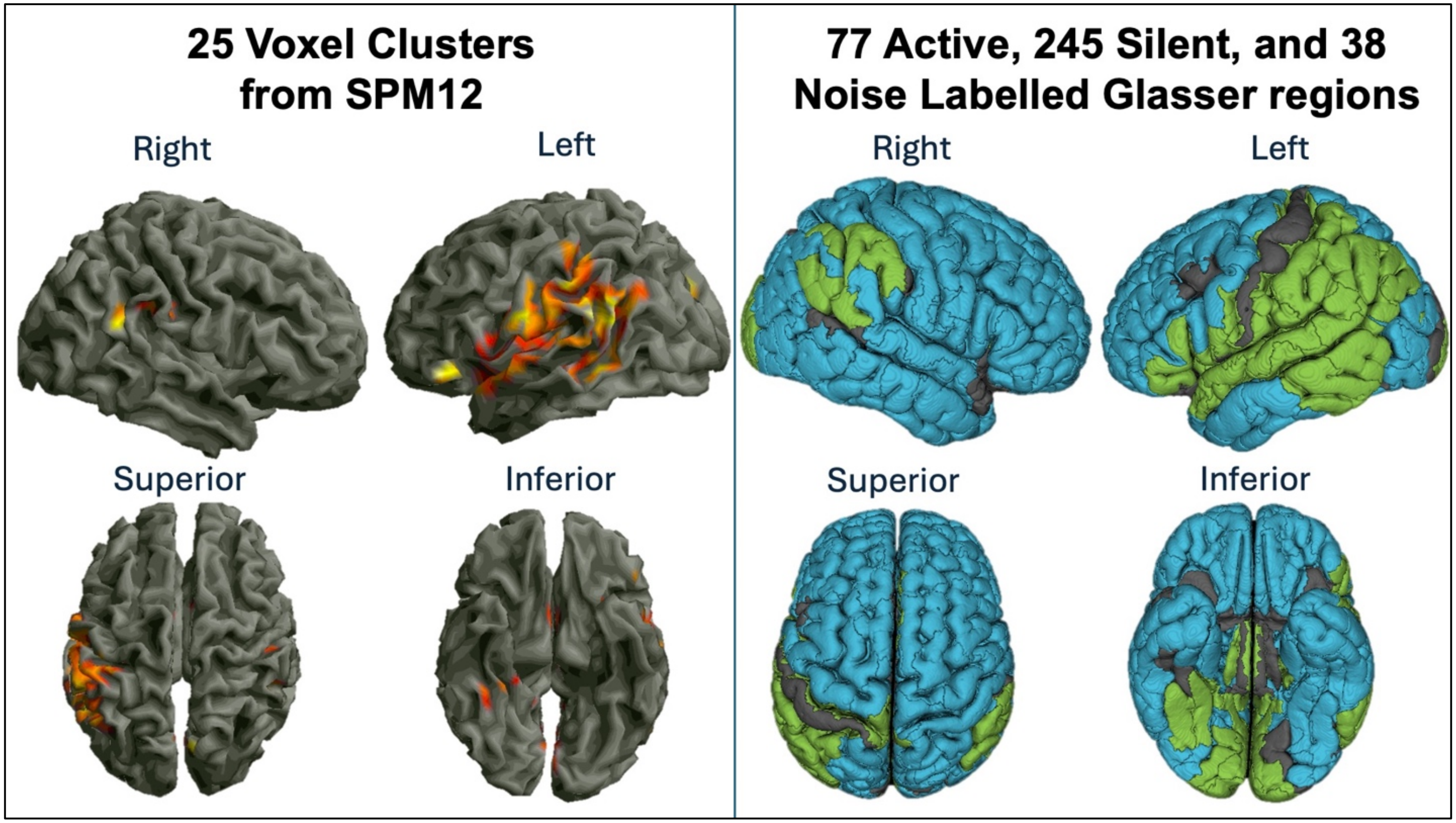

Supplemental Table 3: Correlation of cognitive scores with active regions for the healthy control group. FHR group did not show any significant correlation with cognitive measures. P-values were corrected using FDR correction.

| Active Regions Correlated with Cognitive Measures for Healthy Controls | | | | |
|---|---|---|---|---|
| Region Name | Cognitive Measure | Correlation | df | Corrected p |
| '''R_Area_PGi''' | 0-back response time | 0.612606 | 27 | 0.031695 |
| '''R_Parieto-Occipital_Sulcus_Area_1''' | 2-back response time | 0.719257 | 27 | 0.000847 |
| '''L_Rostral_Area_6''' | 2-back response time | -0.68621 | 27 | 0.000945 |
| '''R_RetroSplenial_Complex''' | 2-back response time | 0.673882 | 27 | 0.000945 |
| '''R_Parieto-Occipital_Sulcus_Area_2''' | 2-back response time | 0.688984 | 27 | 0.000945 |
| '''R_PreSubiculum''' | 2-back response time | 0.679923 | 27 | 0.000945 |
| '''L_Area_TA2''' | 2-back response time | -0.64341 | 27 | 0.001603 |
| '''R_Area_8BM''' | 2-back response time | -0.64719 | 27 | 0.001603 |
| '''R_ParaHippocampal_Area_1''' | 2-back response time | 0.647754 | 27 | 0.001603 |
| '''L_Area_45''' | 2-back response time | -0.62584 | 27 | 0.002415 |
| '''L_Area_ventral_23_a+b''' | 2-back response time | 0.62068 | 27 | 0.002523 |
| '''L_Area_43''' | 2-back response time | -0.61201 | 27 | 0.002686 |
| '''L_Area_IntraParietal_0''' | 2-back response time | 0.614867 | 27 | 0.002686 |
| '''R_Area_7m''' | 2-back response time | 0.603666 | 27 | 0.003116 |
| '''L_RetroSplenial_Complex''' | 2-back response time | 0.593916 | 27 | 0.003573 |
| '''L_Area_Posterior_Insular_1''' | 2-back response time | -0.59067 | 27 | 0.003573 |
| '''R_Area_23c''' | 2-back response time | 0.592162 | 27 | 0.003573 |
| '''L_Area_STSd_anterior''' | 2-back response time | -0.56937 | 27 | 0.005415 |
| '''R_Area_PGi''' | 2-back response time | 0.570304 | 27 | 0.005415 |
| '''L_Area_PGi''' | 2-back response time | 0.565366 | 27 | 0.005649 |
| '''L_Frontal_Opercular_Area_4''' | 2-back response time | -0.53646 | 27 | 0.009898 |
| '''R_Area_Posterior_24_prime''' | 2-back response time | 0.537053 | 27 | 0.009898 |
| '''L_Para-Insular_Area''' | 2-back response time | -0.52066 | 27 | 0.013242 |
| '''L_Auditory_5_Complex''' | 2-back response time | -0.51746 | 27 | 0.013537 |
| '''L_Area_7m''' | 2-back response time | 0.496567 | 27 | 0.019711 |

| | | | | |
|---|---|---|---|---|
| '''L_Area_TemporoParietoOccipital_Junction_2''' | 2-back response time | 0.477504 | 27 | 0.027122 |
| '''R_Superior_Temporal_Visual_Area''' | 2-back response time | 0.470181 | 27 | 0.029791 |
| '''L_Area_31a''' | 2-back response time | 0.456395 | 27 | 0.036578 |
| '''L_Area_STSv_posterior''' | 2-back response time | -0.43923 | 27 | 0.047109 |
| '''R_Area_31a''' | 2-back response time | 0.434555 | 27 | 0.049099 |

Supplemental Table 4: Correlation of cognitive scores with silent regions for the healthy control group. FHR group did not show any significant correlation with cognitive measures. P-values were corrected using FDR correction.

| Silent Regions Correlated with Cognitive Measures for Healthy Controls | | | | |
|---|---|---|---|---|
| Region Name | Cognitive Measure | Correlation | df | Corrected p |
| '''L_Anterior_24_prime''' | 0-back response time | -0.59295 | 27 | 0.017768 |
| '''R_Area_V3B''' | 0-back response time | 0.620998 | 27 | 0.017768 |
| '''R_Primary_Auditory_Cortex''' | 0-back response time | 0.617247 | 27 | 0.017768 |
| '''R_RetroInsular_Cortex''' | 0-back response time | 0.589105 | 27 | 0.017768 |
| '''R_Posterior_Insular_Area_2''' | 0-back response time | 0.592427 | 27 | 0.017768 |
| '''R_Auditory_5_Complex''' | 0-back response time | 0.600196 | 27 | 0.017768 |
| '''R_Area_STSv_posterior''' | 0-back response time | 0.584472 | 27 | 0.017768 |
| '''R_Area_PGp''' | 0-back response time | 0.645851 | 27 | 0.017768 |
| '''R_Area_31pd''' | 0-back response time | 0.588568 | 27 | 0.017768 |
| '''R_Auditory_4_Complex''' | 0-back response time | 0.585956 | 27 | 0.017768 |
| '''R_Area_STSv_anterior''' | 0-back response time | 0.64001 | 27 | 0.017768 |
| '''R_Area_TE1_Middle''' | 0-back response time | 0.587084 | 27 | 0.017768 |
| '''R_ParaHippocampal_Area_3''' | 0-back response time | 0.570609 | 27 | 0.017773 |
| '''R_Area_TemporoParietoOccipital_Junction_3''' | 0-back response time | 0.570444 | 27 | 0.017773 |
| '''R_Area_PGs''' | 0-back response time | 0.572375 | 27 | 0.017773 |
| '''R_Area_FST''' | 0-back response time | 0.575406 | 27 | 0.017773 |
| '''R_Area_posterior_24''' | 0-back response time | 0.571238 | 27 | 0.017773 |
| '''R_IntraParietal_Sulcus_Area_1''' | 0-back response time | 0.561448 | 27 | 0.020823 |
| '''R_Area_STSd_posterior''' | 0-back response time | 0.55701 | 27 | 0.021891 |
| '''R_Area_OP4-PV''' | 0-back response time | 0.543907 | 27 | 0.028049 |
| '''L_Superior_6-8_Transitional_Area''' | 0-back response time | -0.53666 | 27 | 0.030624 |
| '''R_VentroMedial_Visual_Area_1''' | 0-back response time | 0.535618 | 27 | 0.030624 |
| '''L_Area_46''' | 0-back response time | -0.52636 | 27 | 0.032889 |
| '''R_Area_TG_dorsal''' | 0-back response time | 0.529069 | 27 | 0.032889 |
| '''R_Area_Frontal_Opercular''' | 0-back response time | 0.52698 | 27 | 0.032889 |
| '''R_Area_OP1-SII''' | 0-back response time | -0.52368 | 27 | 0.033464 |

| | | | | |
|---|---|---|---|---|
| '''L_Area_44''' | 0-back response time | -0.51511 | 27 | 0.033906 |
| '''L_Area_9-46d''' | 0-back response time | -0.51547 | 27 | 0.033906 |
| '''R_Seventh_Visual_Area''' | 0-back response time | 0.517385 | 27 | 0.033906 |
| '''R_Medial_IntraParietal_Area''' | 0-back response time | 0.517519 | 27 | 0.033906 |
| '''R_Area_OP2-3-VS''' | 0-back response time | 0.514577 | 27 | 0.033906 |
| '''R_Frontal_Opercular_Area_4''' | 0-back response time | 0.511204 | 27 | 0.035183 |
| '''R_Anterior_24_prime''' | 0-back response time | -0.50909 | 27 | 0.035606 |
| '''L_Area_anterior_10p''' | 0-back response time | -0.5031 | 27 | 0.038952 |
| '''R_Area_posterior_10p''' | 0-back response time | 0.500135 | 27 | 0.040113 |
| '''R_Area_V6A''' | 0-back response time | 0.49727 | 27 | 0.04013 |
| '''R_Para-Insular_Area''' | 0-back response time | 0.497839 | 27 | 0.04013 |
| '''R_Area_52''' | 0-back response time | 0.492799 | 27 | 0.042601 |
| '''L_Inferior_6-8_Transitional_Area''' | 0-back response time | -0.4885 | 27 | 0.045059 |
| '''L_Area_dorsal_32''' | 0-back response time | -0.48009 | 27 | 0.045428 |
| '''L_Area_8Av''' | 0-back response time | -0.47903 | 27 | 0.045428 |
| '''L_Area_8Ad''' | 0-back response time | -0.47701 | 27 | 0.045428 |
| '''L_Area_8B_Lateral''' | 0-back response time | -0.48504 | 27 | 0.045428 |
| '''L_Area_anterior_9-46v''' | 0-back response time | -0.48087 | 27 | 0.045428 |
| '''L_Area_posterior_47r''' | 0-back response time | -0.48011 | 27 | 0.045428 |
| '''R_Area_ventral_23_a+b''' | 0-back response time | 0.486485 | 27 | 0.045428 |
| '''R_VentroMedial_Visual_Area_3''' | 0-back response time | 0.478167 | 27 | 0.045428 |
| '''R_Area_Lateral_Occipital_3''' | 0-back response time | 0.476924 | 27 | 0.045428 |
| '''L_Sixth_Visual_Area''' | 0-back response time | 0.471656 | 27 | 0.047432 |
| '''L_Area_11l''' | 0-back response time | -0.47122 | 27 | 0.047432 |
| '''R_Medial_Superior_Temporal_Area''' | 0-back response time | 0.471433 | 27 | 0.047432 |
| '''R_Area_V3B''' | 2-back response time | 0.799475 | 27 | 4.83E-05 |
| '''R_Area_PGs''' | 2-back response time | 0.779256 | 27 | 7.77E-05 |
| '''R_IntraParietal_Sulcus_Area_1''' | 2-back response time | 0.738964 | 27 | 0.00023 |
| '''R_Posterior_Insular_Area_2''' | 2-back response time | 0.739694 | 27 | 0.00023 |
| '''R_Area_PGp''' | 2-back response time | 0.744063 | 27 | 0.00023 |

| | | | | |
|---|---|---|---|---|
| '''R_Area_31pd''' | 2-back response time | 0.723713 | 27 | 0.000373 |
| '''R_Area_STSv_anterior''' | 2-back response time | 0.709248 | 27 | 0.000578 |
| '''R_Medial_IntraParietal_Area''' | 2-back response time | 0.700359 | 27 | 0.000716 |
| '''R_Area_FST''' | 2-back response time | 0.692188 | 27 | 0.000867 |
| '''L_Area_8Ad''' | 2-back response time | -0.67236 | 27 | 0.001108 |
| '''L_Area_44''' | 2-back response time | -0.67855 | 27 | 0.001108 |
| '''L_Superior_6-8_Transitional_Area''' | 2-back response time | -0.67097 | 27 | 0.001108 |
| '''R_RetroInsular_Cortex''' | 2-back response time | 0.67376 | 27 | 0.001108 |
| '''R_ParaHippocampal_Area_3''' | 2-back response time | 0.673917 | 27 | 0.001108 |
| '''R_Area_STSv_posterior''' | 2-back response time | 0.672144 | 27 | 0.001108 |
| '''R_Primary_Auditory_Cortex''' | 2-back response time | 0.665039 | 27 | 0.001195 |
| '''R_Area_ventral_23_a+b''' | 2-back response time | 0.666617 | 27 | 0.001195 |
| '''L_Sixth_Visual_Area''' | 2-back response time | 0.661307 | 27 | 0.00125 |
| '''R_Area_dorsal_32''' | 2-back response time | -0.66034 | 27 | 0.00125 |
| '''R_Seventh_Visual_Area''' | 2-back response time | 0.65876 | 27 | 0.001251 |
| '''L_Area_9-46d''' | 2-back response time | -0.65719 | 27 | 0.001254 |
| '''L_Area_dorsal_32''' | 2-back response time | -0.65352 | 27 | 0.00127 |
| '''L_Area_IFJa''' | 2-back response time | -0.65266 | 27 | 0.00127 |
| '''L_Area_TE1_Middle''' | 2-back response time | -0.65511 | 27 | 0.00127 |
| '''L_Area_8B_Lateral''' | 2-back response time | -0.65098 | 27 | 0.001286 |
| '''L_Area_46''' | 2-back response time | -0.64799 | 27 | 0.001322 |
| '''R_Area_TE1_Middle''' | 2-back response time | 0.647683 | 27 | 0.001322 |
| '''R_Area_OP4-PV''' | 2-back response time | 0.645566 | 27 | 0.001342 |
| '''R_VentroMedial_Visual_Area_3''' | 2-back response time | 0.644915 | 27 | 0.001342 |
| '''R_Auditory_5_Complex''' | 2-back response time | 0.639997 | 27 | 0.001488 |
| '''R_VentroMedial_Visual_Area_1''' | 2-back response time | 0.638909 | 27 | 0.001488 |
| '''R_Auditory_4_Complex''' | 2-back response time | 0.638364 | 27 | 0.001488 |
| '''L_Area_9_Posterior''' | 2-back response time | -0.63269 | 27 | 0.001712 |
| '''L_IntraParietal_Sulcus_Area_1''' | 2-back response time | 0.62903 | 27 | 0.001853 |
| '''R_Area_8Ad''' | 2-back response time | -0.62797 | 27 | 0.001857 |

| | | | | |
|---|---|---|---|---|
| '''L_Anterior_24_prime''' | 2-back response time | -0.62483 | 27 | 0.001978 |
| '''L_Area_IFSp''' | 2-back response time | -0.61973 | 27 | 0.00223 |
| '''R_Area_Lateral_IntraParietal_ventral''' | 2-back response time | 0.615043 | 27 | 0.002479 |
| '''L_Area_8Av''' | 2-back response time | -0.61231 | 27 | 0.002608 |
| '''R_Area_V6A''' | 2-back response time | 0.60865 | 27 | 0.002813 |
| '''R_Area_TemporoParietoOccipital_Junction_3''' | 2-back response time | 0.605488 | 27 | 0.002992 |
| '''L_Dorsal_Transitional_Visual_Area''' | 2-back response time | 0.59884 | 27 | 0.003494 |
| '''L_Area_9_Middle''' | 2-back response time | -0.59779 | 27 | 0.003509 |
| '''R_Area_posterior_24''' | 2-back response time | 0.596111 | 27 | 0.003585 |
| '''L_Seventh_Visual_Area''' | 2-back response time | 0.592816 | 27 | 0.003739 |
| '''R_Area_TG_dorsal''' | 2-back response time | 0.593258 | 27 | 0.003739 |
| '''L_Area_posterior_9-46v''' | 2-back response time | -0.59082 | 27 | 0.003855 |
| '''L_Area__IFSa''' | 2-back response time | -0.58949 | 27 | 0.003906 |
| '''R_Area_9_Posterior''' | 2-back response time | -0.58721 | 27 | 0.004058 |
| '''R_Posterior_InferoTemporal''' | 2-back response time | -0.58476 | 27 | 0.004142 |
| '''R_Ventral_Area_24d''' | 2-back response time | 0.584632 | 27 | 0.004142 |
| '''R_Area_8B_Lateral''' | 2-back response time | -0.58332 | 27 | 0.004142 |
| '''R_Area_s32''' | 2-back response time | 0.583458 | 27 | 0.004142 |
| '''L_Parieto-Occipital_Sulcus_Area_2''' | 2-back response time | 0.574575 | 27 | 0.005057 |
| '''L_Frontal_Eye_Fields''' | 2-back response time | -0.56894 | 27 | 0.005534 |
| '''R_Area_STSd_posterior''' | 2-back response time | 0.568664 | 27 | 0.005534 |
| '''R_Area_OP1-SII''' | 2-back response time | -0.5687 | 27 | 0.005534 |
| '''R_Frontal_Opercular_Area_4''' | 2-back response time | 0.564501 | 27 | 0.00601 |
| '''L_Premotor_Eye_Fields''' | 2-back response time | -0.56018 | 27 | 0.006546 |
| '''R_Area_OP2-3-VS''' | 2-back response time | 0.558827 | 27 | 0.006644 |
| '''L_Area_V3B''' | 2-back response time | 0.557138 | 27 | 0.006751 |
| '''L_Area_anterior_32_prime''' | 2-back response time | -0.55605 | 27 | 0.006751 |
| '''R_Area_9_Middle''' | 2-back response time | -0.55664 | 27 | 0.006751 |
| '''R_Area_Lateral_Occipital_3''' | 2-back response time | 0.551884 | 27 | 0.007316 |
| '''R_Area_46''' | 2-back response time | -0.54967 | 27 | 0.007577 |

| | | | | |
|---|---|---|---|---|
| '''L_Area_TE2_anterior''' | 2-back response time | -0.54632 | 27 | 0.007867 |
| '''L_Area_TemporoParietoOccipital_Junction_3''' | 2-back response time | 0.546025 | 27 | 0.007867 |
| '''L_Area_Lateral_Occipital_3''' | 2-back response time | 0.547213 | 27 | 0.007867 |
| '''R_posterior_OFC_Complex''' | 2-back response time | -0.54412 | 27 | 0.008092 |
| '''L_Area_Lateral_Occipital_2''' | 2-back response time | -0.53932 | 27 | 0.008874 |
| '''R_Area_IntraParietal_0''' | 2-back response time | 0.536646 | 27 | 0.009278 |
| '''R_Area_52''' | 2-back response time | 0.53379 | 27 | 0.009737 |
| '''L_Area_p32_prime''' | 2-back response time | -0.53119 | 27 | 0.010123 |
| '''L_Inferior_6-8_Transitional_Area''' | 2-back response time | -0.53072 | 27 | 0.010123 |
| '''L_Area_25''' | 2-back response time | -0.52769 | 27 | 0.010658 |
| '''L_Area_9_anterior''' | 2-back response time | -0.52643 | 27 | 0.010781 |
| '''R_Medial_Superior_Temporal_Area''' | 2-back response time | 0.525907 | 27 | 0.010781 |
| '''L_Lateral_Area_7P''' | 2-back response time | 0.523575 | 27 | 0.01118 |
| '''L_Area_anterior_9-46v''' | 2-back response time | -0.52247 | 27 | 0.011298 |
| '''L_Area_anterior_10p''' | 2-back response time | -0.51871 | 27 | 0.01202 |
| '''L_Area_TF''' | 2-back response time | -0.51741 | 27 | 0.01202 |
| '''L_Area_STSv_anterior''' | 2-back response time | -0.51711 | 27 | 0.01202 |
| '''R_Dorsal_Area_24d''' | 2-back response time | 0.517719 | 27 | 0.01202 |
| '''R_Area_6_anterior''' | 2-back response time | 0.512952 | 27 | 0.012935 |
| '''L_Area_11l''' | 2-back response time | -0.51079 | 27 | 0.013357 |
| '''L_Area_31pd''' | 2-back response time | 0.509733 | 27 | 0.013487 |
| '''L_Ventral_Area_6''' | 2-back response time | -0.50829 | 27 | 0.013569 |
| '''R_Area_Frontal_Opercular''' | 2-back response time | 0.508729 | 27 | 0.013569 |
| '''L_Area_55b''' | 2-back response time | -0.5059 | 27 | 0.014076 |
| '''R_Anterior_24_prime''' | 2-back response time | -0.50525 | 27 | 0.014098 |
| '''L_Area_TE1_anterior''' | 2-back response time | -0.50269 | 27 | 0.014672 |
| '''R_Area_posterior_10p''' | 2-back response time | 0.502114 | 27 | 0.014678 |
| '''L_VentroMedial_Visual_Area_2''' | 2-back response time | -0.50103 | 27 | 0.014834 |
| '''L_Medial_Area_7P''' | 2-back response time | 0.499082 | 27 | 0.015247 |
| '''L_Area_V3A''' | 2-back response time | 0.490519 | 27 | 0.0178 |

| | | | | |
|---|---|---|---|---|
| '''R_Area_V3A''' | 2-back response time | 0.489171 | 27 | 0.018073 |
| '''R_Entorhinal_Cortex''' | 2-back response time | 0.487987 | 27 | 0.018106 |
| '''R_Area_PF_opercular''' | 2-back response time | 0.488498 | 27 | 0.018106 |
| '''R_Area_43''' | 2-back response time | 0.485755 | 27 | 0.018694 |
| '''R_Area_9-46d''' | 2-back response time | -0.48325 | 27 | 0.019396 |
| '''L_Area_posterior_10p''' | 2-back response time | -0.47946 | 27 | 0.02045 |
| '''R_Area_anterior_32_prime''' | 2-back response time | 0.479335 | 27 | 0.02045 |
| '''L_Area_dorsal_23_a+b''' | 2-back response time | 0.478428 | 27 | 0.02049 |
| '''L_Area_31p_ventral''' | 2-back response time | 0.478177 | 27 | 0.02049 |
| '''L_Polar_10p''' | 2-back response time | -0.47176 | 27 | 0.022596 |
| '''L_Area_PGp''' | 2-back response time | 0.472131 | 27 | 0.022596 |
| '''L_Ventral_Area_24d''' | 2-back response time | 0.466029 | 27 | 0.024807 |
| '''L_Medial_Area_7A''' | 2-back response time | 0.463079 | 27 | 0.025894 |
| '''L_Area_posterior_47r''' | 2-back response time | -0.46204 | 27 | 0.026129 |
| '''R_Area_31p_ventral''' | 2-back response time | 0.454022 | 27 | 0.029509 |
| '''R_Area_TG_Ventral''' | 2-back response time | 0.453987 | 27 | 0.029509 |
| '''R_Area_5L''' | 2-back response time | 0.451222 | 27 | 0.03066 |
| '''R_Lateral_Area_7P''' | 2-back response time | 0.449996 | 27 | 0.031029 |
| '''R_Area_TemporoParietoOccipital_Junction_2''' | 2-back response time | 0.447861 | 27 | 0.031888 |
| '''R_Medial_Area_7P''' | 2-back response time | 0.446292 | 27 | 0.032456 |
| '''R_Ventral_Area_6''' | 2-back response time | 0.444646 | 27 | 0.033076 |
| '''L_Parieto-Occipital_Sulcus_Area_1''' | 2-back response time | 0.441633 | 27 | 0.0343 |
| '''R_Area_Posterior_Insular_1''' | 2-back response time | 0.441433 | 27 | 0.0343 |
| '''R_Middle_Temporal_Area''' | 2-back response time | 0.439169 | 27 | 0.035306 |
| '''R_Area_8C''' | 2-back response time | -0.43351 | 27 | 0.0384 |
| '''R_VentroMedial_Visual_Area_2''' | 2-back response time | 0.430577 | 27 | 0.039926 |
| '''L_Area_TG_Ventral''' | 2-back response time | -0.42978 | 27 | 0.040106 |
| '''R_Area_5m_ventral''' | 2-back response time | 0.424573 | 27 | 0.043216 |
| '''R_Area_2''' | 2-back response time | 0.42359 | 27 | 0.043536 |
| '''L_Ventral_Visual_Complex''' | 2-back response time | 0.422388 | 27 | 0.044012 |

| | | | | |
|---|---|---|---|---|
| '''R_Area_TA2''' | 2-back response time | 0.415829 | 27 | 0.048347 |
| '''L_Area_6m_anterior''' | 2-back response time | -0.41484 | 27 | 0.048704 |
| '''R_Para-Insular_Area''' | 2-back response time | 0.41401 | 27 | 0.048939 |
| '''R_Area_dorsal_23_a+b''' | 2-back response time | 0.412598 | 27 | 0.049238 |
| '''R_Area_STSd_anterior''' | 2-back response time | 0.413002 | 27 | 0.049238 |

Supplemental Table 5: Average regional betweenness and eigenvector centrality for active and silent networks for both FHR and control groups. Average and standard deviation across groups is shown. Statistical comparisons between active and silent networks across FHR and control groups for both average betweenness and eigenvector centrality. The p-values are Bonferroni corrected. ***Between group comparisons of graph metrics of active and silent networks were not significant; therefore, they are not included in the table.**

| Comparison of Undirected Graph Measure between Active and Silent Networks within groups | | | | | | | | |
|---|---|---|---|---|---|---|---|---|
| | HC | | | | FHR | | | |
| **Measure** | **Active** | **Silent** | **F (df)** | **p** | **Active** | **Silent** | **F(df)** | **p** |
| Betweenness Centrality | 77.93±29.6 | 268.5±75.8 | 19.02 | 0.002 | 87.05±39.0 | 326.9±89.3 | 0.20 | >0.05 |
| Eigenvector Centrality | 0.093±0.011 | 0.049±0.0049 | 43.63 | <0.001 | 0.092±0.010 | 0.043±0.0042 | 1.21 | >0.05 |
| Clustering Coefficient | 0.69±0.096 | 0.65±0.078 | 49.04 | <0.001 | 0.67±0.083 | 0.65±0.067 | 4.30 | >0.05 |
| Degree | 25.89±12.1 | 79.93±43.6 | 13.64 | 0.018 | 23.55±11.5 | 84.05±34.1 | 0.05 | >0.05 |

Supplemental Table 6: Group-wise average of global metrics. Mean and standard deviation across the group is shown for all global graph metrics calculated.

| Group-wise average of global metrics | | | | |
|---|---|---|---|---|
| | HC | | FHR | |
| **Measure** | **Active** | **Silent** | **Active** | **Silent** |
| Assortativity | 0.272±0.16 | 0.399±0.13 | 0.315±0.15 | 0.363±0.09 |
| Pathlength | 1.99±0.39 | 2.09±0.33 | 2.08±0.52 | 2.10±0.37 |
| Modularity | 0.302±0.14 | 0.302±0.12 | 0.282±0.15 | 0.273±0.11 |

**Hub and Module Analysis:**

Full network: In the full network of 360 regions, we identified 18 hubs. Of the 18, 6 were common between groups which were part of visual, motor, and inferior frontal cortex (**Supplemental Table 7**). Hubs unique to FHR were in the anterior cingulate and medial prefrontal cortex, but the HC had medial temporal, orbital and polar frontal, and dorsolateral prefrontal cortex regions.

We examined the modules for consistency by running the code 10 times. FHR full network consisted of 8 modules whereas the HC had 7. Each module had at least one hub except 2 modules for FHR and 1 for control that did not comprise of a hub.

Active network: FHR and HC hubs were unique to each group (**Supplemental Table 8**). Hubs in the HC AN consisted of lateral temporal and auditory association cortex whereas FHR hubs consisted of primary visual, posterior cingulate, and Temporo-Parieto-Occipital (TPO) Junction.

Both groups had two modules and hubs were present in all modules. The regions assigned to each module were similar between groups with only 3.9% difference (3 regions assigned to different modules). These regions are the left Area 22 prime (anterior cingulate and medial prefrontal cortex), left Area PFm Complex (Inferior Parietal Cortex), and left Superior Temporal Visual Area (TPO Junction).

Silent network: The HC and FHR SN had 3 common hubs which were located in the paracentral lobule and mid cingulate, orbital and polar frontal, and premotor cortexes (**Supplemental Table 9**). For FHR, the unique hubs consisted of inferior frontal, insular and frontal opercular, anterior cingulate and medial prefrontal, somatosensory and motor, early auditory, and inferior frontal cortexes. HC hubs were in MT complex and neighboring visual areas, early visual, ventral stream visual, medial temporal, dorsolateral prefrontal, and posterior opercular cortex.

The silent networks had 6 modules for FHR and 3 modules for HC. No hubs were in two of the smallest modules for the FHR whereas each module consisted of at least two hubs for the HC.

Supplemental Table 7: Hubs in the full networks of the FHR and controls. The Glasser section and the module number the hub was located in is shown. Red font indicates overlapping hubs between the control and FHR groups.

| | Healthy Controls | | | | Familial High-Risk | | | |
|---|---|---|---|---|---|---|---|---|
| **Rank** | **Region Name** | **Section** | **S=0, A=1, N=3** | **Module** | **Region Name** | **Section** | **S=0, A=1, N=3** | **Module** |
| 1 | L Primary Visual Cortex | Primary Visual Cortex | 1 | 2 | L Primary Visual Cortex | Primary Visual Cortex | 1 | 6 |
| 2 | R Second Visual Area | Early Visual Cortex | 1 | 2 | L Area 3a | Somatosensory and Motor Cortex | 1 | 4 |
| 3 | R Second Visual Area | Early Visual Cortex | 1 | 2 | R Area 44 | Inferior Frontal Cortex | 0 | 7 |
| 4 | L Area 3a | Somatosensory and Motor Cortex | 1 | 1 | L Second Visual Area | Early Visual Cortex | 1 | 6 |
| 5 | L Primary Motor Cortex | Somatosensory and Motor Cortex | 1 | 1 | L Primary Motor Cortex | Somatosensory and Motor Cortex | 1 | 4 |
| 6 | R ParaHippocampal Area 3 | Medial Temporal Cortex | 0 | 2 | L Area STSv posterior | Auditory Association Cortex | 1 | 4 |
| 7 | L Area 23c | Paracentral Lobular and Mid Cingulate Cortex | 3 | 3 | L Area PHT | Lateral Temporal Cortex | 1 | 2 |
| 8 | L Area 11l | Orbital and Polar Frontal Cortex | 0 | 7 | L Area posterior 24 | Anterior Cingulate and Medial Prefrontal Cortex | 0 | 3 |
| 9 | R Area 5m ventral | Paracentral Lobular and Mid Cingulate Cortex | 0 | 3 | R Second Visual Area | Early Visual Cortex | 1 | 6 |
| 10 | R Auditory 5 Complex | Auditory Association Cortex | 0 | 5 | R Supplementary and Cingulate Eye Field | Paracentral Lobular and Mid Cingulate Cortex | 0 | 8 |
| 11 | R Area PHT | Lateral Temporal Cortex | 0 | 2 | R Area TE1 posterior | Lateral Temporal Cortex | 0 | 6 |
| 12 | R Area V4t | MT Complex and Neighboring Visual Areas | 0 | 2 | L Third Visual Area | Early Visual Cortex | 3 | 6 |
| 13 | L Third Visual Area | Early Visual Cortex | 3 | 2 | L Supplementary and Cingulate Eye Field | Paracentral Lobular and Mid Cingulate Cortex | 0 | 4 |
| 14 | R Third Visual Area | Early Visual Cortex | 1 | 2 | L Anterior 24 prime | Anterior Cingulate and Medial Prefrontal Cortex | 0 | 3 |
| 15 | R Superior Frontal Language Area | Dorsolateral Prefrontal Cortex | 0 | 7 | L Rostral Area 6 | Premotor Cortex | 1 | 4 |
| 16 | R Area 44 | Inferior Frontal Cortex | 0 | 6 | L Area 6 anterior | Premotor Cortex | 0 | 4 |
| 17 | R Rostral Area 6 | Premotor Cortex | 0 | 5 | L Area V4t | MT Complex and Neighboring Visual Areas | 0 | 6 |
| 18 | R Area 11l | Orbital and Polar Frontal Cortex | 0 | 7 | L Area STSv anterior | Auditory Association Cortex | 0 | 4 |

Supplemental Table 8: Hubs in the active networks of the FHR and controls. The Glasser section and the module number the hub was located in is shown. Red font indicates overlapping hubs between the control and FHR groups.

| | Healthy Controls | | | | Familial High Risk | | | |
|---|---|---|---|---|---|---|---|---|
| **Rank** | **Region Name** | **Section Label** | **S=0, A=1, N=3** | **Module** | **Region Name** | **Section Label** | **S=0, A=1, N=3** | **Module** |
| 1 | L Area PHT | Lateral Temporal Cortex | 1 | 1 | L Primary Visual Cortex | Primary Visual Cortex | 1 | 2 |
| 2 | L Area 3a | Somatosensory and Motor Cortex | 1 | 2 | L Primary Motor Cortex | Somatosensory and Motor Cortex | 1 | 1 |
| 3 | L Area TA2 | Auditory Association Cortex | 1 | 2 | L RetroSplenial Complex | Posterior Cingulate Cortex | 1 | 2 |
| 4 | L Area STGa | Auditory Association Cortex | 1 | 2 | L Area TemporoParietoOccipital Junction 2 | Temporo-Parieto-Occipital Junction | 1 | 2 |

Supplemental Table 9: Hubs in the silent networks of the FHR and controls. The Glasser section and the module number the hub was located in is shown. Red font indicates overlapping hubs between the control and FHR groups.

| | Healthy Controls | | | | Familial High Risk | | | |
|---|---|---|---|---|---|---|---|---|
| **Rank** | **Region Name** | **Section Label** | **S=0, A=1, N=3** | **Module** | **Region Name** | **Section Label** | **S=0, A=1, N=3** | **Module** |
| 1 | R Area 5m ventral | Paracentral Lobular and Mid Cingulate Cortex | 0 | 1 | R Area 44 | Inferior Frontal Cortex | 0 | 1 |
| 2 | R Area V4t | MT Complex and Neighboring Visual Areas | 0 | 1 | R Area Posterior Insular 1 | Insular and Frontal Opercular Cortex | 0 | 1 |
| 3 | L Fourth Visual Area | Early Visual Cortex | 0 | 1 | R Area 13l | Orbital and Polar Frontal Cortex | 0 | 5 |
| 4 | L Supplementary and Cingulate Eye Field | Paracentral Lobular and Mid Cingulate Cortex | 0 | 2 | R Area posterior 24 | Anterior Cingulate and Medial Prefrontal Cortex | 0 | 5 |
| 5 | L Area 11l | Orbital and Polar Frontal Cortex | 0 | 2 | R Area TE1 posterior | Lateral Temporal Cortex | 0 | 6 |
| 6 | L Ventral Visual Complex | Ventral Stream Visual Cortex | 0 | 1 | R Primary Motor Cortex | Somatosensory and Motor Cortex | 0 | 1 |
| 7 | R ParaHippocampal Area 3 | Medial Temporal Cortex | 0 | 1 | R Primary Sensory Cortex | Somatosensory and Motor Cortex | 0 | 1 |
| 8 | R Area PHT | Lateral Temporal Cortex | 0 | 1 | R Area 2 | Somatosensory and Motor Cortex | 0 | 1 |
| 9 | R Area 8Av | Dorsolateral Prefrontal Cortex | 0 | 2 | R Rostral Area 6 | Premotor Cortex | 0 | 1 |
| 10 | R Area 13l | Orbital and Polar Frontal Cortex | 0 | 2 | R RetroInsular Cortex | Early Auditory Cortex | 0 | 1 |
| 11 | R Rostral Area 6 | Premotor Cortex | 0 | 3 | L Supplementary and Cingulate Eye Field | Paracentral Lobular and Mid Cingulate Cortex | 0 | 2 |
| 12 | R Area OP2-3/VS | Posterior Opercular Cortex | 0 | 3 | L Area 44 | Inferior Frontal Cortex | 0 | 2 |
